# Hemodynamics in Patients with Aortic Coarctation: A Comparison of *in vivo* 4D-Flow MRI and FSI Simulation

**DOI:** 10.1101/2023.02.13.528355

**Authors:** Priya J. Nair, Martin R. Pfaller, Seraina A. Dual, Michael Loecher, Doff B. McElhinney, Daniel B. Ennis, Alison L. Marsden

## Abstract

The analysis of quantitative hemodynamics provides information for the diagnosis and treatment planning in patients with aortic coarctation (CoA). Patient-specific computational fluid dynamics (CFD) simulations reveal detailed hemodynamic information, but their agreement with the clinical standard 4D-Flow magnetic resonance imaging (MRI) needs to be characterized. This work directly compares *in vivo* CFD fluid-structure interaction (FSI) simulations against 4D-Flow MRI in patients with CoA (N=5). 4D-Flow MRI-derived flow waveforms and cuff blood pressure measurements were used to tune the boundary conditions for the FSI simulations. Flow rates from 4D-Flow MRI and FSI were compared at cross-sections in the ascending aorta (AAo), CoA and descending aorta (DAo). Qualitative comparisons showed an overall agreement of flow patterns in the aorta between the two methods. The *R*^2^ values for the flow waveforms in the AAo, CoA, and DAo were 0.97, 0.84 and 0.81 respectively, representing a strong correlation between 4DFlow MRI measurements and FSI results. This work characterizes the use of patient-specific FSI simulations in quantifying and analyzing CoA hemodynamics to inform CoA treatment planning.

## 1 Introduction

Coarctation of the aorta (CoA) is a congenital heart defect characterized by a segmental constriction of the aorta. It accounts for 6-8% of all congenital heart defects [7], with an estimated incidence of 3 per 10,000 live births [8]. The narrowing of the aorta causes a drop in blood pressure (BP) across the CoA. If left untreated, it can result in hypertension, stroke, and aortic rupture [1]. Even after successful repair (defined as correction of the BP drop), patients with CoA continue to experience long-term ventricular and arterial complications, and have significantly lower survival rates than the normal population [3]. Previous studies have related the morbidity in CoA to adverse local hemodynamics in the aorta and its branches [4]. Studying and understanding the hemodynamics associated with CoA and its progression is therefore of high clinical importance.

CFD has proved to be a useful tool for the assessment of local hemodynamics in patients with cardiovascular disease. Non-invasive anatomic and physiological data acquired in the clinic can be used to generate patient-specific hemodynamics simulations to inform diagnosis, treatment and outcomes. While CFD approaches show great potential, the method must be characterized against the clinical standard: *in vivo* 4D-Flow MRI. In this study, we compare qualitative and quantitative hemodynamics from patient-specific CFD FSI simulations to those obtained from 4D-Flow MRI in five patients with CoA.

### 2 Methods

### 2.1 CFD FSI Simulation

#### Patient Data Acquisition

4D-Flow MRI datasets (IRB approved) were acquired for five patients with CoA (Table 1) who thereafter underwent a catheter-based stenting procedure. Cuff BP was measured non-invasively on the same day. Imaging exams were not conducted on the day of the catheterization but were acquired, on average, 80 days (range = 4-222 days) before catheterization. Average difference in heart rate between the day of imaging and the day of catheterization was 3 bpm, indicating similar physiological states on both days.

**Table 1.** Patient characteristics.

| Patient | Sex | Age (y) | BSA (m<sup>2</sup>) | Cuff BP (mmHg) systolic/diastolic | CO (L/min) | HR (bpm) |
| --- | --- | --- | --- | --- | --- | --- |
| P-1 | Male | 15 | 1.94 | 145/86 | 8.9 | 68 |
| P-2 | Female | 54 | 1.56 | 154/54 | 3.8 | 67 |
| P-3 | Male | 21 | 1.58 | 105/61 | 3.3 | 94 |
| P-4 | Male | 22 | 1.89 | 123/47 | 8.4 | 63 |
| P-5 | Male | 44 | 2.09 | 152/93 | 6.4 | 54 |

#### Anatomic Model Generation

3D geometries of patient-specific aortas were generated by importing the MRI magnitude images into SimVascular and manually segmenting the vessels. Centerlines were automatically extracted from the 3D geometry to create the 0D model comprising a network of lumped parameter elements (resistors, capacitors and inductors) [5]. The 3D volume was discretized using a mesh of tetrahedral elements. A three-layer boundary mesh was incorporated to resolve the velocity gradients at the wall. The mesh was further refined at the CoA and the post-stenotic dilation region to capture the jet formed by the CoA. The final mesh had approximately two million tetrahedral elements.

#### Boundary Conditions

Boundary conditions were tuned using non-invasive cuff BP measurements, and flow measurements derived from 4D-Flow MRI. Eddy current-corrected 4D-Flow MRI datasets were used to measure 2D time-resolved flow at the inlets and outlets using Arterys (Arterys, San Francisco, USA). 0D simulations were run first and used to tune the boundary conditions. The patientspecific temporally varying flow profile was prescribed to the 0D model inlet. Flow splits to each of the aortic branches, along with cuff BP data, were used to tune outflow boundary conditions. Three-element Windkessel boundary conditions (proximal resistance R_*p*_, capacitance C, distal resistance R_*d*_) were imposed at the outlets. Total resistance (R_*p*_ + R_*d*_) for each of the branches was calculated using the cuff BP and the flow split determined from MRI. C as well as the R_*p*_/R_*d*_ ratio were adjusted to fine-tune the boundary conditions until the calculated pressures matched the patient’s systolic and diastolic cuff BP within 5 mmHg. The tuned boundary conditions were then applied to the 3D model outlets (Figure 1). The patient-specific flow waveform derived from 4D-Flow was prescribed to the 3D model inlet, assuming a parabolic flow profile.

**Fig. 1.**
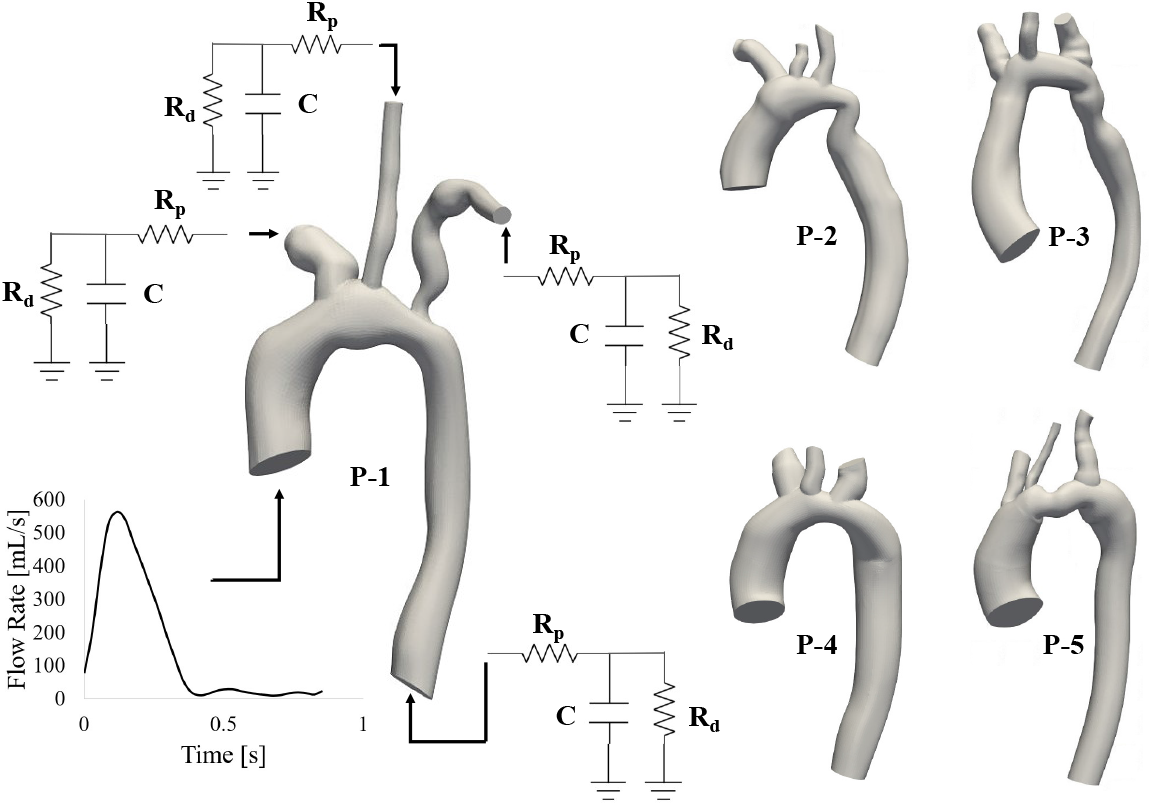
3D patient-specific anatomic models generated by segmenting MRI images, with depiction of boundary conditions applied. R_*p*_: proximal resistance, C: capacitance, R_*d*_: distal resistance

#### Simulations

Simulations were performed with svSolver, SimVascular’s finite element solver for fluid-structure interaction between an incompressible, Newtonian fluid and a linear elastic membrane for the vascular wall. The Young’s modulus of the aortic wall was defined to be 3×10^6^ dyn/cm^2^, based on previously reported values of stiffness in a human aorta with CoA [2]. The wall thickness was assumed to be 10% of the vessel diameter. After optimizing the boundary conditions, the 0D simulation was run first for 10 cardiac cycles. The results from the last cardiac cycle of the 0D simulation were projected onto the 3D mesh and used to initialize the 3D simulation, allowing the 3D simulation to converge to a steady solution faster than it would with zero or steady state initialization [6].

The 3D simulations were performed with a deformable wall using the coupled momentum method (CMM). The model was prestressed by applying a diastolic load. For the 3D simulations, Poisson’s ratio = 0.5 and density of blood = 1 g/cm^3^ were used. The 3D simulations were run for 10 cardiac cycles to ensure convergence; only the last cardiac cycle was analyzed. Time-resolved velocity was extracted from the last simulation cycle. At each centerline point, time-resolved cross-sectional average velocities were computed. FSI results had a temporal resolution of 1 ms and were visualized using ParaView.

### 2.2 4D-Flow MRI

Patient-specific 4D-Flow datasets were accessible through Arterys. Eddy current correction was applied using a machine learning-based correction tool available in Arterys and then manually checked. Aorta model geometry from a single timepoint was used to mask 4D-Flow velocity fields. The dicom images were clipped to include only regions that had non-zero velocity and image magnitude. The resulting velocity fields were averaged over the cross-section using the same method described for averaging FSI results and were visualized using ParaView.

### 2.3 Statistical Analysis

Errors were computed as the difference between FSI and 4D-Flow-derived measurements, then averaged across patients. *R*^2^ values of flow waveforms from FSI and 4D-Flow MRI at slices in the AAo, CoA and DAo were computed.

## 3 Results

Qualitative comparisons of flow patterns reveal a good match between the *in vivo* 4D-Flow MRI-derived velocity fields and those obtained from the FSI simulation at peak systole (Figure 2). At the same time, the velocity magnitude of the 4DFlow data sometimes appears substantially lower than that of FSI.

**Fig. 2.**
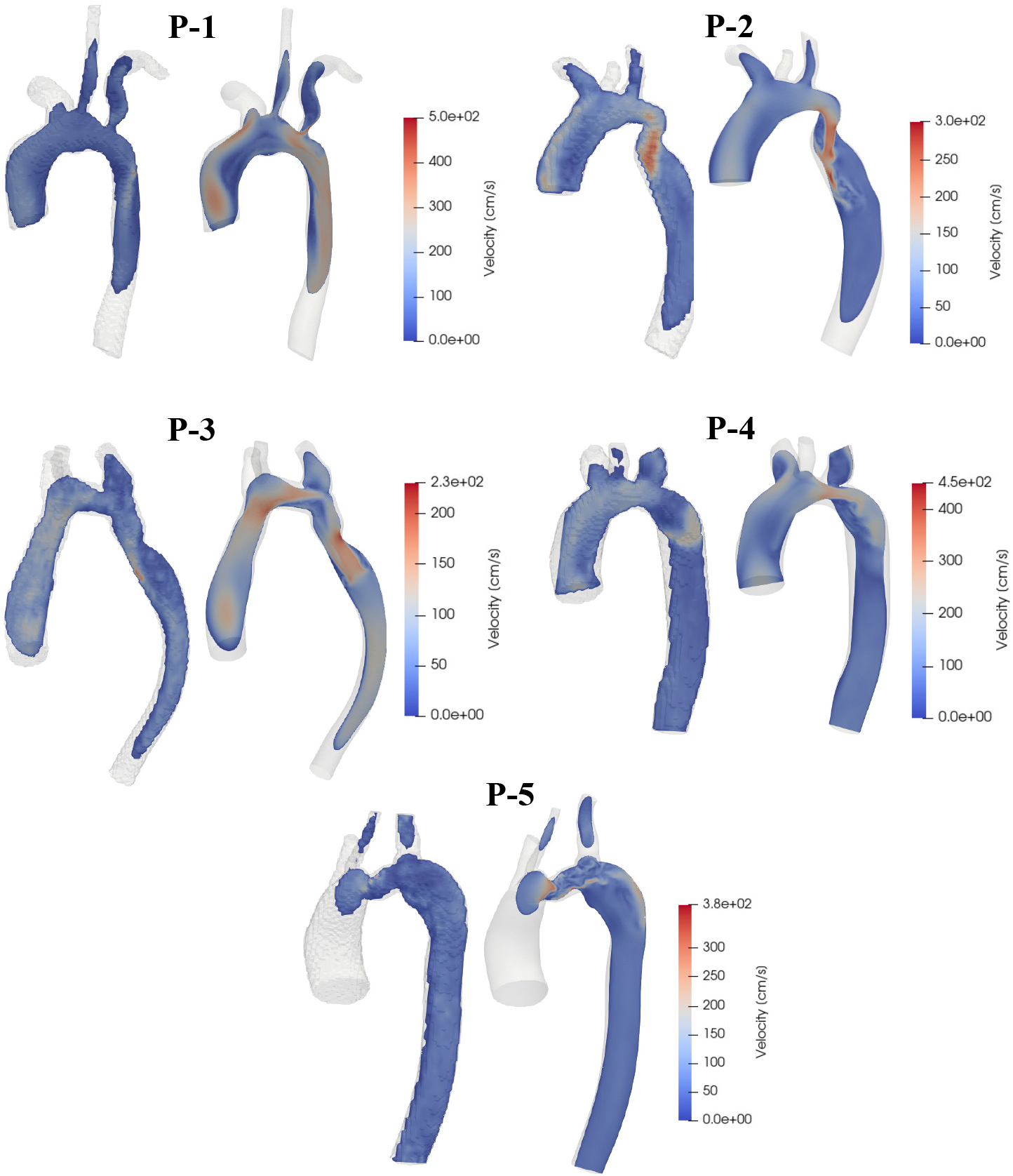
Velocity magnitude at peak systole on a sagittal plane, a comparison between *in vivo* 4D-Flow MRI (left) and FSI simulation (right).

Quantitative comparisons of flow rate at the AAo, CoA and DAo slice-levels show a general agreement of flow waveforms between 4D-Flow MRI and FSI simulations (Figure 3). In P-1 and P-3, a time delay was observed between the occurrence of peak flow in 4D-Flow versus FSI in the CoA and DAo. A lack of conservation of mass in the 4D-Flow MRI flow measurements was also observed, with the mean flow in the DAo exceeding the CoA flow in 3 out of 5 patients. FSI tends to overestimate the mean flow rate, peak flow rate, and peak mean velocity in all slices. The error was minimum at the AAo, where the 4DFlow MRI-derived flow waveform was prescribed as a boundary condition, and maximum at the CoA (Table 3). Errors in mean flow rate from FSI in the AAo, CoA and DAo were 13.5%, 85.4% and 29% of the mean flow rate measured using 4D-Flow MRI. Similarly, errors in peak flow rate were 10.7%, 68.6% and 37.8%. Errors in peak mean velocity were 18.8%, 88.6% and 53.4%. *R*^2^ for the flow waveforms in the AAo, CoA, and DAo were 0.97, 0.84 and 0.81 respectively. This represents a strong correlation between 4D-Flow measurements and FSI results, with decreasing goodness of fit as distance from the inlet increases.

**Fig. 3.**
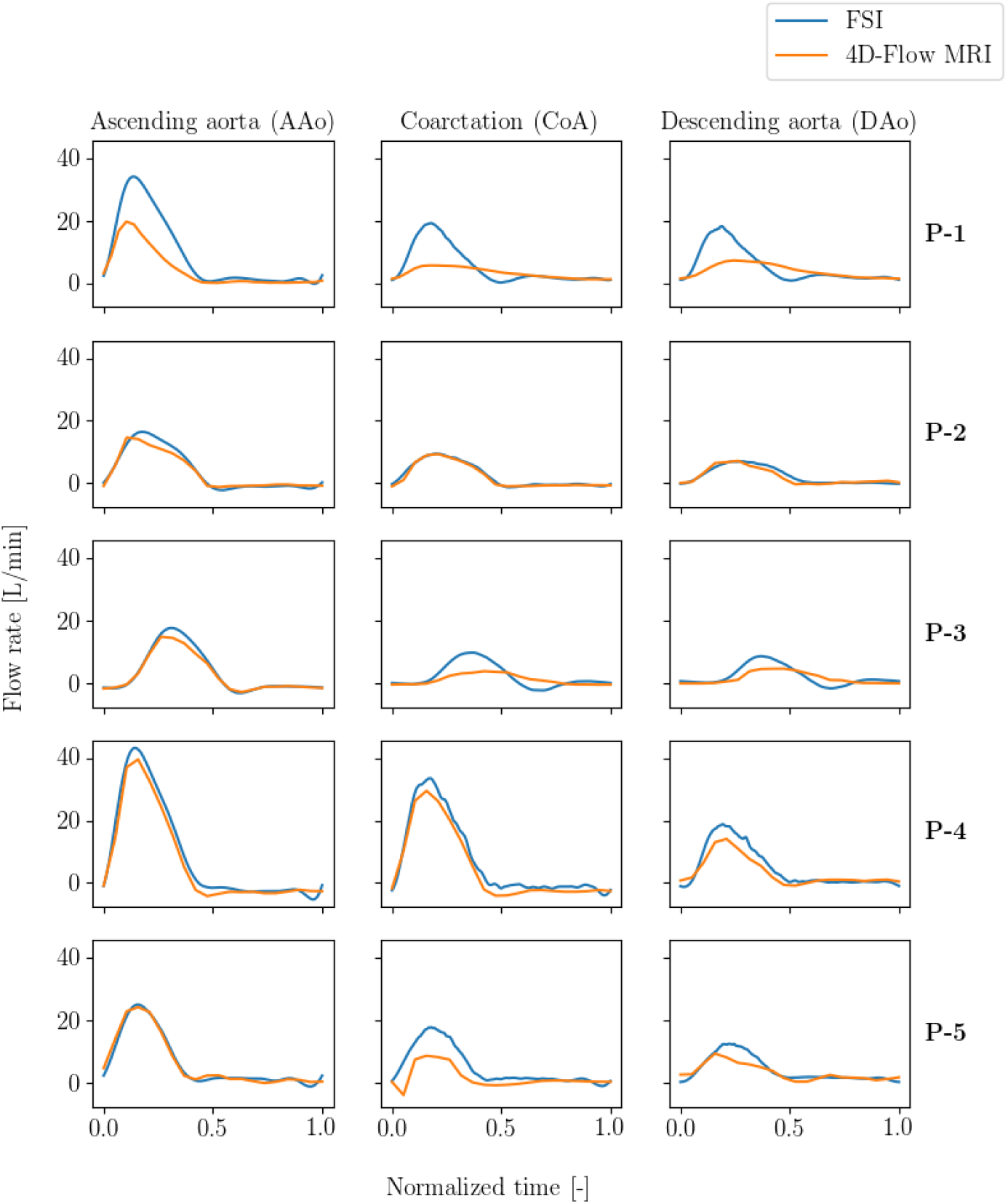
Comparison of *in vivo* 4D-Flow MRI and simulated volumetric flow rate over time at a slice each in the AAo, CoA, and DAo.

**Table 2.** 4D-Flow MRI imaging parameters.

| Patient | VENC (cm/s) | Temporal resolution (ms) | Pixel size (mm<sup>2</sup>) |
| --- | --- | --- | --- |
| P-1 | 350 | 28.4 | 1.2 x 1.2 |
| P-2 | 550 | 42.5 | 1.4 x 1.8 |
| P-3 | 250 | 30.3 | 0.9 x 1.3 |
| P-4 | 300 | 42.3 | 1.3 x 1.8 |
| P-5 | 350 | 52.8 | 1.4 x 1.9 |

**Table 3.** Flow characteristics.

| Patient | Location | Mean flow rate<br>(L/min) |  | Peak flow rate<br>(L/min) |  | Peak mean<br>velocity (cm/s) |  |
| --- | --- | --- | --- | --- | --- | --- | --- |
|  |  | 4D | FSI | 4D | FSI | 4D | FSI |
| P-1* | AAo | 4.5 | 9.1 | 19.7 | 34.2 | 98.4 | 185.0 |
|  | CoA | 3.3 | 5.4 | 5.7 | 19.3 | 63.3 | 240.0 |
|  | DAo | 4.0 | 5.4 | 7.3 | 18.3 | 68.3 | 193.4 |
| P-2 | AAo | 3.6 | 4.0 | 14.6 | 16.4 | 50.0 | 58.3 |
|  | CoA | 2.2 | 2.4 | 9.3 | 9.4 | 101.7 | 131.7 |
|  | DAo | 2.1 | 2.4 | 7.2 | 7.0 | 41.7 | 45.0 |
| P-3 | AAo | 2.9 | 2.5 | 14.9 | 17.6 | 56.7 | 71.7 |
|  | CoA | 1.1 | 2.2 | 3.8 | 9.7 | 56.7 | 140.0 |
|  | DAo | 1.6 | 2.2 | 4.6 | 8.6 | 46.7 | 98.4 |
| P-4 | AAo | 7.1 | 8.8 | 39.7 | 43.3 | 98.4 | 116.7 |
|  | CoA | 5.2 | 7.2 | 29.6 | 33.6 | 98.4 | 121.7 |
|  | DAo | 3.5 | 4.8 | 14.1 | 18.9 | 60.0 | 88.4 |
| P-5 | AAo | 6.6 | 6.5 | 24.3 | 25.1 | 36.7 | 41.7 |
|  | CoA | 1.7 | 5.0 | 8.7 | 17.8 | 80.0 | 203.4 |
|  | DAo | 3.3 | 4.2 | 9.4 | 12.5 | 38.3 | 56.7 |
| **Error** | AAo | 0.65 ± 0.78 | | 2.22 ± 1.20 | | 11.67 ± 6.09 | |
| | CoA | 1.65 ± 1.32 | | 4.78 ± 3.76 | | 65.01 ± 47.27 | |
| | DAo | 0.78 ± 0.43 | | 2.92 ± 2.20 | | 25.42 ± 20.30 | |

\*Excluded from summary statistics

In P-1 specifically, large errors were observed between 4D-Flow MRI and FSI results, which may be explained by the observation that the vessel significantly expanded/contracted over the cardiac cycle. This large wall deformation caused discrepancies in the measured hemodynamic quantities given the limits of our current vessel wall registration process. Results from P-1 although reported in the study, are therefore excluded from the summary statistics.

## 4 Discussion

This study aimed to characterize the agreement of FSI simulations with the clinical standard 4D-Flow MRI in five patients with CoA. Qualitative comparisons between 4D-Flow and FSI showed that patient-specific simulations are capable of capturing the overall flow patterns seen in patients with CoA.

Quantitative FSI results showed discrepancies in mean flow rate, peak flow rate, and peak mean velocity when compared to 4D-Flow. Some of the errors can be explained by the uncertainties associated with the 4D-Flow MRI technique.

The sub-optimal spatial and temporal resolution of images acquired using 4DFlow can contribute to discrepancies we see in the flow fields. Additionally, while several corrections are applied to the acquired velocity field data, some errors from phase offsets may still be present. This can be confirmed by our observation of lack of conservation of mass in the flow measured using 4D-Flow MRI in this study. Some of the errors observed in this study can also be attributed to the registration process used to mask the 4D-Flow MRI velocity data. We used the aortic geometry from a single time point to mask velocity information acquired throughout the cardiac cycle. For patients whose vessels experience large deformations over the cardiac cycle, our current registration process would introduce discrepancies in flow measurements.

On the FSI side, we used the same vessel wall stiffness (from literature) for all five patients. The mismatch of this defined stiffness with the true stiffness of the patient’s aorta may explain the mismatch of the flow waveforms seen at the CoA and DAo. This likely also contributed to the time delay in the flow rate peak between 4D-Flow and FSI in P-1 and P-3. Lastly, the use of parabolic velocity profiles at the inlet could also have resulted in discrepancies downstream. Use of a more comprehensive registration process and methods to determine patientspecific stiffness estimates for the vessel walls can be explored in the future.

Overall, this study characterizes the hemodynamic similarities and differences between 4D-Flow MRI and FSI simulations. It establishes FSI simulations as a valuable alternative to long MRI scans in assessing hemodynamics to inform treatment planning in patients with CoA.

